# Origin of imported SARS-CoV-2 strains in The Gambia identified from whole genome sequences

**DOI:** 10.1101/2020.10.26.354969

**Authors:** Abdoulie Kanteh, Jarra Manneh, Sona Jabang, Mariama A. Kujabi, Bakary Sanyang, Mary A. Oboh, Abdoulie Bojang, Haruna S. Jallow, Davis Nwakanma, Ousman Secka, Anna Roca, Alfred Amambua-Ngwa, Martin Antonio, Ignatius Baldeh, Karen Forrest, Ahmadou Lamin Samateh, Umberto D’Alessandro, Abdul Karim Sesay

## Abstract

Severe Acute Respiratory Syndrome Coronavirus 2 (SARS-CoV-2) is a positive-sense single stranded RNA virus with high human transmissibility. This study generated Whole Genome data to determine the origin and pattern of transmission of SARS-CoV-2 from the first six cases tested in The Gambia. Total RNA from SARS-CoV-2 was extracted from inactivated nasopharyngeal-oropharyngeal swabs of six cases and converted to cDNA following the ARTIC COVID-19 sequencing protocol. Libraries were constructed with the NEBNext ultra II DNA library prep kit for Illumina and Oxford Nanopore Ligation sequencing kit and sequenced on Illumina MiSeq and Nanopore GridION, respectively. Sequencing reads were mapped to the Wuhan reference genome and compared to eleven other SARS-CoV-2 strains of Asian, European and American origins. A phylogenetic tree was constructed with the consensus genomes for local and non-African strains. Three of the Gambian strains had a European origin (UK and Spain), two strains were of Asian origin (Japan). In The Gambia, Nanopore and Illumina sequencers were successfully used to identify the sources of SARS-CoV-2 infection in COVID-19 cases.

## Introduction

The emerging and re-emerging of pathogens such as severe acute respiratory syndrome-coronavirus 2 (SARS-CoV-2) pose a grave threat to human health.^1^ The SARS-CoV-2 disease, first detected in Wuhan, China, in December 2019 has become a global pandemic,^2^ and is causing an unprecedented burden on the health care systems and economies globally.^3–6^ Worldwide, the number of cases has been increasing exponentially,^6^ especially in Europe and America, with significant but variable case-fatality rates between continents. By April 28^th^, 2020, there were more than 3.1 million SARS-CoV-2 confirmed cases and more than 200,000 deaths.^7^ Nevertheless, SARS-CoV-2 confirmed cases in sub-Saharan Africa are currently relatively low, possibly due to much lower international air traffic than in other continents and thus a low number of imported cases.^8^ By the 28^th^ April 2020, The Gambia, a tourism hotspot, had reported a total of ten SARS-CoV-2 cases, including one death. While the travel history of index cases may suggest the origin of infection, phylogenetic analysis of the strains isolated from these cases and contacts will provide a precise link between local transmission and other global populations.

The first SARS-CoV-2 case was reported to be an acquired zoonotic infection,^9,10^ followed by efficient and rapid human-to-human transmission from Wuhan, China, to other Asian countries and then other continents.^11–13^ The single stranded positive sense RNA genome of the SARS-CoV-2 is closely related to the Middle East Respiratory Syndrome-Coronavirus (MERS-CoV) and the Severe Acute Respiratory Syndrome Coronavirus (SARS-CoV).^9–14^ These pathogens pose significant risk to global health and modern-day life, hence the need for effective strategies to detect the sources of infections, outbreaks and transmission patterns in different geographical settings.

The phylogenetic analyses of global SARS-CoV-2 sequences provide insight into the relatedness of strains from different areas and suggest the transmission of four super-clades,^15^ geographically clustering into viral isolates from Asia (China), US (two super clades) and Europe. The objective of this analysis was to provide genome data on six cases of SARS-CoV-2 in The Gambia, determine the source of these strains, baseline for subsequent local transmission, and contribute genomic diversity data towards local and global vaccine design. The Oxford Nanopore GridION and Illumina MiSeq platforms were utilized to sequence the viral genomes from four confirmed SARS-CoV-2, one inconclusive and one negative case by rRT-PCR. We also analysed the genomes of samples classified as indeterminate and negative by RT-PCR (COVID-19 detection assay) from two different cases respectively.

## Methods

### Sample acquisition

Nasopharyngeal-Oropharyngeal (NP-OP) swabs (451) from SARS-CoV-2 suspected cases and their contacts were transported to the Medical Research Council Unit The Gambia at London School of Hygiene and Tropical Medicine (MRCG at LSHTM). Of the 451 samples screened by rRT-PCR, ten were confirmed as SARS-CoV-2 cases and 5 as indeterminate cases (positive for the screening gene and negative for the SARS-CoV-2 confirmatory gene)

For WGS, four SARS-CoV-2 confirmed cases, one indeterminate case and one negative case were processed Table 1. In one of the confirmed cases, different isolates from samples collected up to 10 days apart were sequenced. Of the 6 cases sequenced, 4 were male; 2 females, there was one death, two recoveries and two active cases.

**Table 1:** Sample information for COVID-19 sequenced cases from The Gambia

| Case ID | Travelled from | Date Reported | Current Status | Number of samples submitted | Time points | Library prep type |  |  | Sequencing |  |
| --- | --- | --- | --- | --- | --- | --- | --- | --- | --- | --- |
|  |  |  |  |  |  | Depletion | ARTIC amplicon (NEB) | ARTIC amplicon (ONT -LSK109) | Illumina (MiSeq) | Nanopore (GridION) |
| A | London | 16/03/20 | Recovered | 4 | Days 0,4,7,10 | 2 | 4 | 4 | 4 | 4 |
| B | Bangladesh | 19/03/20 | Dead | 1 | Day 0 | 0 | 1 | 1 | 1 | 1 |
| C | France | 20/03/20 | Recovered | 1 | Day 0, | 0 | 1 | 1 | 1 | 1 |
| D | France | 26/03/20 | Active | 2 | Day 0,11 | 0 | 1 | 2 | 1 | 2 |
| E | Netherland | 23/03/20 | Active | 2 | Day 0,14 | 0 | 1 | 2 | 1 | 2 |
| F | Italy | 13/03/20 | Recovered | 1 | Day 0 | 0 | 1 | 1 | 1 | 1 |
| Total |  |  |  | 11 |  | 2 | 9 | 11 | 9 | 11 |
| Cases A-D = Confirmed RT-PCR COVID-19 cases |  |  |  |  |  |  |  |  |  |  |
| Case E = Indeterminate by RT-PCR |  |  |  |  |  |  |  |  |  |  |
| Case F = RT-PCR COVID-19 negative |  |  |  |  |  |  |  |  |  |  |
| Cases A and D travelled to The Gambia in the same flight |  |  |  |  |  |  |  |  |  |  |
| Cases C and D both travelled from France |  |  |  |  |  |  |  |  |  |  |

### RNA extraction

Total RNA was purified from eleven samples Table 1 using the QiaAmp viral RNA mini kit (Qiagen – 52906) following viral inactivation at the MRCG at LSHTM containment level 3 facility. The purified RNA samples were quantified using Qubit RNA reagent kit on a Qubit fluorometer 3.0 (concentration range 3-7 ng/μl) (Invitrogen). RNA integrity (RINe) was checked on the Agilent TapeStation

4200 yielding a RINe range of 2.1–5.

### Ribosomal RNA depletion, cDNA synthesis and Multiplex PCR

Two of the samples (day 0 and 4) from Case A were depleted using the RiboMinus transcription isolation kit from ThermoFisher and purified using RNA purification beads from Beckman Coulter. The purified rRNA-depleted samples were converted to cDNA as per the NEBNext ultra II RNA library prep kit for Illumina (NEB, E7770L). Total RNA from the rest of the samples was converted to cDNA according to the ARTIC amplicon sequencing protocol for SARS-CoV-2.^artic^ ARTIC protocol primer ^artic^ schemes for SARS-CoV-2 (Version 2) were used for the multiplex PCR. Two primer pools at 10 μM containing 98 primers each were used for the PCR amplification. The samples were subjected to 35 cycles of PCR. The purified products were visualised and quantified.

### Illumina and Nanopore library Preparation and Sequencing Illumina and Nanopore library preparation

The purified cDNA from the depletion and PCR products from the ARTIC protocol were normalised to 100 ng with EB buffer (10 mM Tris-HCl) to a final volume of 25 μl for Illumina library preparation using the NEBNext ultra II DNA library prep kit for Illumina (New England Biolabs, UK; E7645). Following 7 cycles of PCR enrichment, the libraries were purified and quantified using the high sensitivity dsDNA Qubit kit and sized using D1000 ScreenTape on the Agilent TapeStation 4200 (amplicon size range 519-572 bp). Each sample was normalised to 10 nM before pooling. The pool was run at a final concentration of 10 pM on an Illumina MiSeq instrument using MiSeq V3 reagent kit. The pool was denatured with sodium hydroxide according to Illumina recommendation and spiked with 5% PhiX (PhiX control v3 Illumina Catalogue FC-110-3001) before loading (Fig 1).

**Fig 1.**
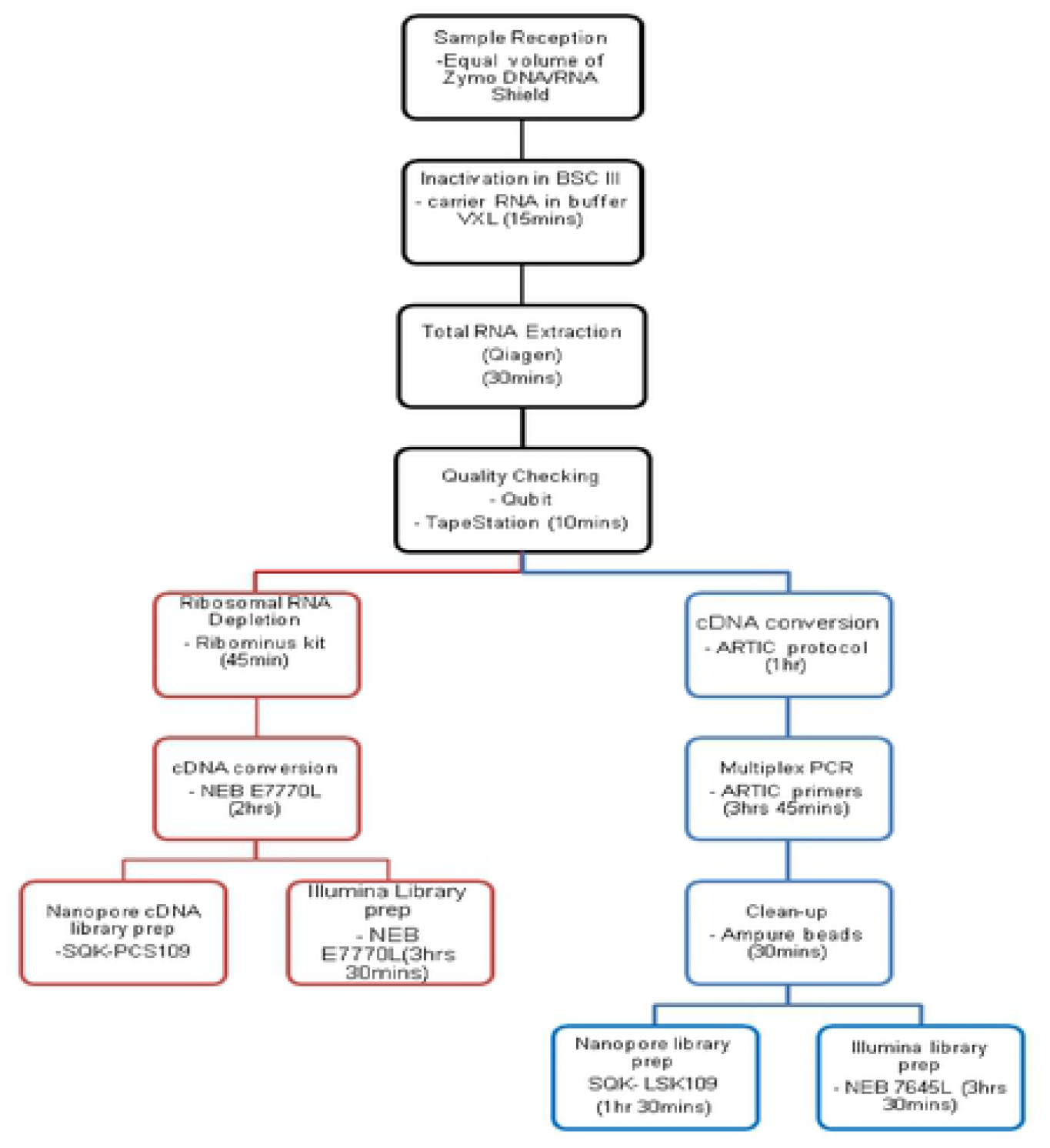
Summary of the Library preparation steps for Illumina and Oxford Nanopore Sequencing Technology platforms. Library preparation took ~ 8 hours for the Nanopore workflow and ~10 hours for the Illumina workflow.

Nanopore sequencing library preparation was performed according to the manufacturer’s instructions for the Ligation Sequencing Kit (SQK-LSK109, Oxford Nanopore Technologies). Briefly, the cDNA samples were amplified using the ARTIC protocol and purified with 1X AMPure XP beads. Individual samples were then subjected to end repair and adapter ligation following SQK-LSK109 protocol. 20 ng of each library was loaded on the Oxford Nanopore GridION on individual R9.4.1 flow cells and sequencing data monitored on the fly using Rampart (v1.1.0). Fig 1 summarised the steps involved in sequencing using for both Nanopore and Illumina platform.

### Quality control and read mapping for Illumina and Nanopore platform

Although a minimum read depth of 30X for the SARS-CoV-2 genome was targeted, more than 100X coverage was generated on both platforms. FASTQ files were subjected to various quality control checks and analysed following standard analysis pipelines (SARS-CoV-2 novel Coronavirus bioinformatics protocol; SAMTOOLS).

For Nanopore data, sequencing reads were quality checked using MinIONQC,^16^ and only reads with a minimum Q score of 7 were included in our subsequent analysis. Quality checked reads were run through what’s in My Pot (WIMP) pipeline on the Oxford Nanopore EPI2ME platform to verify the number of reads characterised as SARS-CoV-2.

We used SARS-CoV-2 novel Coronavirus bioinformatics protocol developed by Nick Loman *et. al*. to analyse the Nanopore data.^17^ Firstly, we used “ARTIC Guppylex” to remove chimeric reads from each sample with the following parameters (--min-length 400 −max-length 700). Filtered reads were then mapped to Wuhan-Hu-1 reference genomes (accession number MN908947.3) and Single nucleotide polymorphism (SNPs) were generated based on the reference and a consensus genome for each sample was generated using SAMTOOLS (v1.9). To further validate our results, we ran all genomes through the coronavirus typing tool (v1.13).

Similarly, 250 bp paired end Illumina reads were quality checked using FASTQC (v0.11.5).^18^ Reads with only a minimum Phred score of 30 were included in our downstream analysis. One sample which was SARS-CoV-2 negative by reverse-transcription real-time polymerase chain reaction (rRT-PCR) was characterised as Bat coronavirus using kraken and thus excluded from the analysis. Filtered reads were then mapped to the Wuhan-Hu-1 reference genomes (accession number MN908947.3) using BWA-mem (0.7.17-r1188). BCFtools Mpileup (v1.8) was used in creating a variant file. Finally, BCFtools consensus was used in generating the FASTA consensus sequence for each sample.

### Phylogenetic analysis for Illumina and Nanopore platform

Prank (v140603) was used to generate a multiple alignment of all the samples including some available reference genomes around the globe (Downloaded from RefSeq). These strains were selected based on the patients’ travel history and the major geographical spread of the pandemic. We finally constructed a maximum likelihood phylogenetic tree using the General time reversible model (GTR) with IQTREE (v1.3.11.1). The Interactive Tree of Life (ITOL) (v5) was used to visualise and annotate the phylogenetic tree.

## Results and Discussion

Whole genome sequencing data was generated from six confirmed cases from both sequencing platforms; the additional time points from cases D and E were sequenced only on the Nanopore GridION (Table 2). Two samples from the first case were sequenced on both platforms following ribosomal depletion, the results generated (not included) showed depletion of human sequences and the majority of the reads mapped to bacterial sequences with only 0.03% from the Illumina reads mapping to the SARS-CoV-2 reference strain. The rRT-PCR and the sequencing data generated are summarized in Table 2.

**Table 2:** Summary of COVID-19 results

| Case ID | Time point | Diagnostic results Covid-19 (Ct) |  | Sequencing results Covid-19 |  | Phylogenetic Inferences |
| --- | --- | --- | --- | --- | --- | --- |
|  |  | Gene 1 (E gene) | Gene 2 (RdRP) | Illumina | Nanopore | Illumina & Nanopore |
| **A** | **Day 0** | + | + | + | + | **Europe (UK)** |
|  | **Day 4** | + | + | + |  |  |
|  | **Day 7** | + | + | + |  |  |
|  | **Day 10** | + | + | + |  |  |
| **B** | **Day 0** | + | + | + |  | **Asia (Japan)** |
| **C** | **Day 0** | + | + | + |  | **Europe (Spain)** |
| **D** | **Day 0** | + | + | + |  | **Europe (Spain)** |
|  | **Day 11** | + | + | *Not sequenced* |  |  |
| **E** | **Day 0** | + | - | + |  | **Asia (Japan)** |
|  | **Day 14** | + | - | *Not sequenced* |  |  |
| **F** | **Day 0** | - | - | - | - |  |

From the Illumina platform, a high quality read length of 250 bp paired-end reads was generated for each sample after 48 hrs post library preparation. Total number of sequences ranged from three to six million reads with an average mapping quality of 60 when mapped to the Wuhan reference genome. The Nanopore platform generated a read length of 400 - 17000 bp for each sample after 12 hrs of sequencing. It recorded a range of one to four million reads with average mapping quality of 58 across the reference genome for each sample. To compare the results from both technologies, consensus genomes were generated for each sample and a maximum likelihood phylogenetic tree was constructed. Both platforms showed a similar topology with 1000 bootstrap clustering The Gambian isolates with the European and Asian strains as illustrated in Figs 2 and 3.

**Figure 2.**
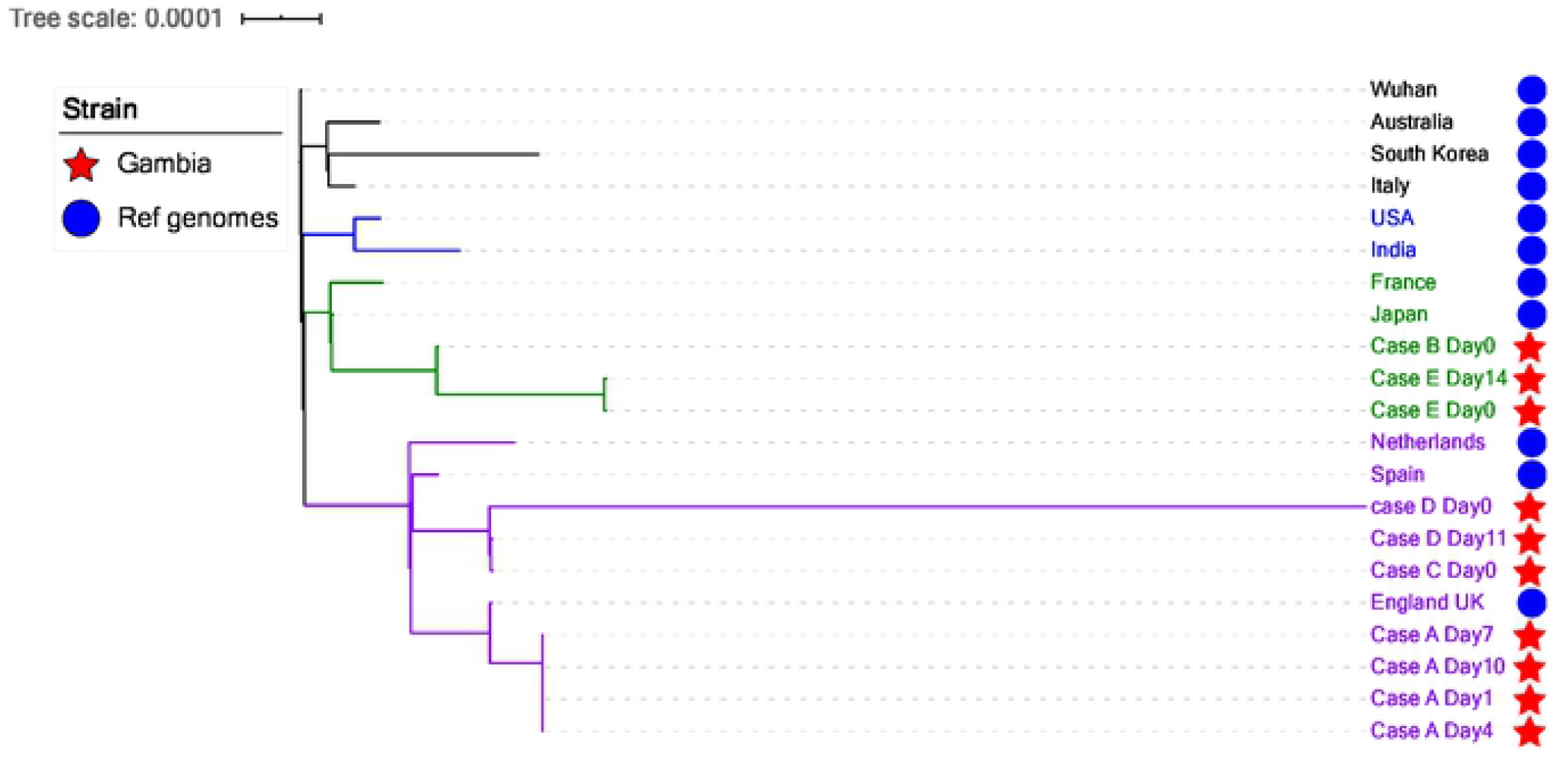
A maximum likelihood phylogenetic tree of ten SARS-CoV-2 genomes isolated from The Gambia (Nanopore data) and 11 SARS-CoV-2 strains isolated in different parts of the world. The tree showed the genetic relation of strains isolated in The Gambian to the global circulating strains.

**Figure 3.**
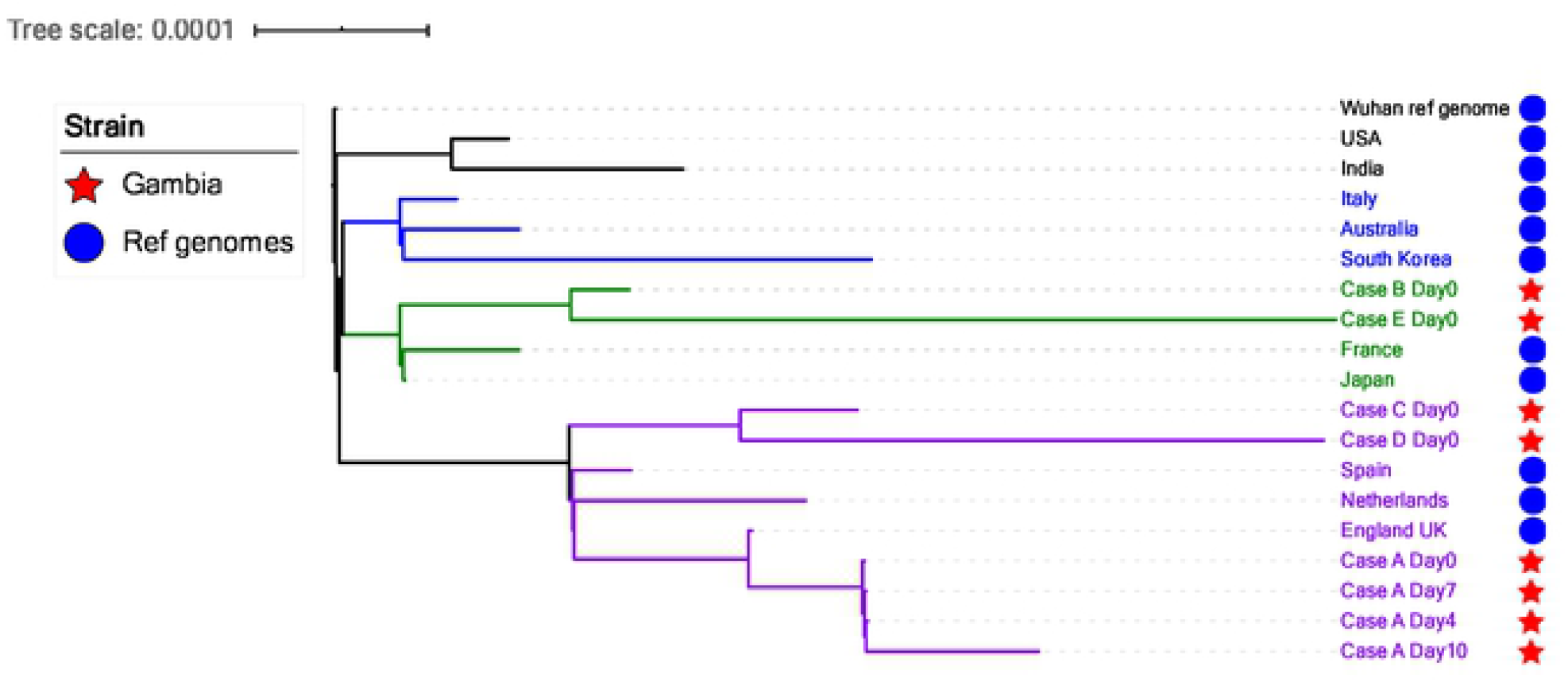
A maximum likelihood phylogenetic tree of eight SARS-CoV-2 genomes isolated from The Gambia (Illumina data) and 11 SARS-CoV-2 strains isolated in different parts of the world. The tree showed the genetic relation of strains isolated in The Gambian to the global circulating strains.

Six genomes (4 samples from Case A, 1 from case D and 1 from case C) from the Gambian samples clustered with the European (Spanish and United Kingdom) SARS-CoV-2 strains. This is not surprising given that these patients had been in Europe before arriving in The Gambia. Although viruses are known to mutate and change rapidly,^19,20^ the viral genome of case A clustered on the same node at different time points indicating the patient was shedding the same virus with no observed polymorphism according to Nanopore results. Interestingly, the same samples sequenced on the MiSeq suggested polymorphism at day 10, resulting in a longer branch length compared to previous time points (Figure 3). Further analysis of the Illumina data showed seven more SNPs in the day 10 sample compared to the other time points. The SNP winked by the Nanopore phylogenetically, might have an associated higher error rate compared to the Illumina. Strains from cases C and D, both having travelled from France, were more closely related to the Spanish strain included for comparison. Though cases D and A travelled to The Gambia on the same flight, their strains had a different origin, indicating that they could have been infected independently, before the start of their journeys.

The viral genome from case B who initiated travel from Bangladesh and then across four other countries, including Senegal, before arriving in The Gambia, clustered with a strain from Japan. This case may have contracted the infection in Asia and his travel history suggests he could have contributed to infections in other countries. The two isolates from case E at different time points clustered with strains from Japan as well. Interestingly, case E samples were indeterminate by rRT-PCR diagnostics, even though the outcome from multiple alignment showed no mismatch between the sequences and the primer set. The indeterminate diagnostic rRT-PCR result could be due to low sensitivity of the assay, an indication of low viral density of SARS-CoV-2 in the sample. Therefore, subsequent follow up for such cases is essential to further evaluate diagnosis and aid towards the understanding of the disease progression and the evolution of this novel virus strain under different case management environments.

Although WGS data is still limited in sub-Saharan Africa, this approach has proven to be a highly sensitive, specific and confirmatory tool for SARS-CoV-2 detection. Hence, the use of second and third generation sequencing technologies coupled with bioinformatics is quite imperative in providing data for monitoring transmission dynamics.

From the two sequencing platforms, we were able to rapidly generate sequencing data in 20 hours and 3 days after sample reception on the Nanopore and Illumina platforms respectively. While Illumina sequencing may be more accurate in determining within-sample-diversity, Nanopore data can help with the understanding of the linkage between SNPs within individual virions. The Nanopore platform with its flexibility for number of samples per run, and the generation of data in real-time and at a reasonable cost makes it most suitable for outbreaks. Therefore, with our optimised and ready-to-go workflow, we are set to generate data for tracking SARS-CoV-2 in The Gambia and other African countries within 24 hours of sample reception. This would go a long way in providing knowledge on the molecular epidemiology of this disease, give the true burden of the disease in this setting (as seen in the resolution of the indeterminate cases) as well as provide information for African specific vaccine development and inform policy makers on decisions for strategic control measures.

## Conclusion

We have demonstrated that the Nanopore platform with the flexibility of high-end desktop sequencer (GridION) to the portable sequencer (MinION) in combination with the ARTIC protocol and workflow allows for cost-effective (wide range for the number of runs and samples per flow cell), and near real-time generation of pathogen sequence data. Our analysis has shown that the SARS-CoV-2 strains identified in The Gambia are of European and Asian origin and sequenced data matched patients’ travel history. In addition, we were able to show that two COVID19 positive cases travelling in the same flight had in fact different sources of infection.

## Acknowledgment

We acknowledge the use of CLIMB server for the cloud-based analysis, the field sample collection by the teams at Ministry of Health, Epidemiology Department, Thushan de Silva for helpful discussion on ARTIC protocol and sequencing, Covid-19 laboratory diagnostic staff, and at MRCG at LSHTM Logistics, Staff at CSD, COVID-19 Emergency Management.

## Authors contribution

AK lead with Nanopore platform and the bioinformatics analysis, JM lead with Illumina platform, MAK lead with viral Inactivation and purification. AK, JM, MAK, SJ, BS, MAO & AB contributed to the sequencing pipeline and writing of the manuscript.

## Competing Interests

The authors declare that they have no competing interests.

The Genomic Core facility at MRCG at LSHTM is the one and only certified service provider for the ONT GridION platform in Africa.

### Ethics statement

The study has under gone under MRCG at LSTM and Gambia Government joint ethics committee for review.

## Data and Materials Availability

The details of methods used in the paper is available as a supplementary document. GISAID submission number: EPI_ISL_428856 and EPI_ISL_428857

The data from the genomes sequenced in the Gambia were submitted and available in Nextstrain website for real-time tracking of the pathogen evolution.

